# Propofol Treatment Alters the Metabolic State of MDA-MB-231 Breast Cancer Cells

**DOI:** 10.1101/2025.01.15.633222

**Authors:** Tasbassum Ahmed Tasmi, Shubha Holla, Alex J. Walsh

## Abstract

The mechanisms of many anesthetic drugs, such as propofol, are unknown. This study investigates the impact of propofol, a widely used intravenous anesthetic drug, on the metabolic behavior of the human triple-negative breast cancer cell line, MDA-MB 231. Utilizing fluorescence lifetime imaging microscopy (FLIM), we assessed the effect of propofol on cellular metabolism through imaging of the fluorescence lifetimes of NADH and FAD, coenzymes of metabolism reactions. Exposure to propofol-induced significant morphological and metabolic changes of the cells, including substantial cellular shrinkage, which may reveal a metabolic component to propofol’s mechanism.

## Introduction

Anesthesia drugs play a crucial role in modern medical procedures, yet significant gaps remain in our understanding of their precise mechanisms of action. While their primary effects on consciousness and pain perception are well-established, the potential impact of these drugs on cellular metabolism is less clear (1). Recent studies suggest that certain anesthetics, including propofol, may influence key metabolic pathways such as mitochondrial function, oxidative stress regulation, and energy production (2). These interactions could have profound implications, especially in pathologies like cancer, where metabolic reprogramming is a hallmark of disease progression (3). Propofol, a known intravenous anesthetic drug, alters cellular metabolism by interacting with mitochondrial function. It can suppress mitochondrial function oxygen consumption, leading to a metabolic shift from oxidative phosphorylation to glycolysis. Such metabolic changes may not only affect normal cellular activities but also impact diseases like cancer, where metabolic reprogramming supports tumor growth and survival (4).

Anesthetic drugs, particularly propofol, have gained attention for potential roles beyond routine anesthesia during surgery. Recent studies have suggested that propofol may have effects on tumor cell behavior by impacting cellular metabolism, proliferation, and invasion (9). Moreover, propofol promotes cell apoptosis while increasing sensitivity to chemotherapeutic drugs (9). Researchers are paying more attention to the intricate relationships that propofol has with cancer cells as they investigate its possible anticancer mechanism.

Breast cancer remains one of the leading causes of cancer-related morbidity and mortality for women globally, emphasizing the urgent need for novel therapeutic strategies (5). Currently, the majority of cancer-related fatalities among American women are caused by breast cancer, which is also the second most common cancer overall (6). Cancer cells alter their metabolic pathways to sustain their unregulated growth and metastasis (7). This shift sustains extensive cell growth and replication. The MDA- MB 231 cell line, often characterized by its aggressive nature and its triple-negative phenotype, serves as a standard model for studying breast cancer (10). It lacks the presence of estrogen receptor (ER), progesterone receptor (PR), and human epidermal growth receptor 2 (HER2), making it resistant to most targeted therapies (11).

Anesthetic drugs, traditionally used for inducing anesthesia, have shown potential altering cancer cell metabolism, which could be leveraged for cancer treatment. These drugs can regulate signaling pathways, induce epigenetic modifications, and interact with chemotherapeutic agents to inhibit tumor growth and enhance treatment efficacy (12) Propofol has been noted for its ability to suppress cancer cell proliferation and invasion (4), while local anesthetics can reduce metastasis by modulating cytoskeletal structures and ion channels.

Fluorescence Lifetime Imaging Microscopy (FLIM) is a potent technique for examining these interactions at the cellular level providing a non-invasive, spatially resolved assessment of metabolic patterns in living cancer cells. Subtle changes in the cellular metabolism can be detected by FLIM through the analysis of NADH and FAD lifetimes as both of these are the indicators of metabolic activity and cellular redox state (13). This capability is especially valuable when studying the effects of treatments on cancer cells, as it allows to observe real-time metabolic responses at the cellular level (14).

Fluorescence Lifetime Imaging Microscopy (FLIM), a non-invasive and spatially resolved label-free imaging technique that measures the fluorescence decay of fluorophores and potentially can be used to measure the metabolic profile of live cancer cells (15). FLIM is a core biomedical imaging tool providing high-resolution images of molecular contrast in living samples. It utilizes the lifetime property of fluorescence and has high sensitivity to the molecular environment and changes in molecular conformation. FLIM has been widely implemented in autofluorescence imaging to investigate cellular metabolism (8). The endogenous fluorescence of NAD(P)H (reduced form of nicotinamide adenine dinucleotide phosphate) and FAD (oxidized form of flavin adenine dinucleotide), two essential coenzymes participating in various critical metabolic pathways, has been used as the optical contrast source in metabolic imaging techniques (16). The fluorescence decay rates depend on whether the coenzymes are free or (protein) bound (17). NAD(P)H has a short lifetime in the free state and a longer lifetime in the bound condition (18) while FAD has a short lifetime in the bound configuration and a longer lifetime in the free state (19) (20).

Our study focuses on understanding if anesthetic drugs affect cellular metabolism. We aimed to elucidate the metabolic consequences of propofol treatment in MDA MB 231 cells by applying fluorescence lifetime imaging microscopy (FLIM). FLIM allows real-time monitoring of metabolic states through the analysis of NADH and FAD lifetimes (21).

## Materials and Methods

### Cell culture and preparation

Triple-negative breast cancer cells, MDA-MB-231, were cultured in high glucose Dulbecco’s Modified Eagle Medium (DMEM) (11965-084), supplemented with 10% Fatal Bovine Serum (FBS) (35-011-CV) and 1% antibiotic-antimycotic (30-004-CI). For fluorescence lifetime imaging, cells were seeded at a density of 1×10^6^ per 35mm glass-bottom imaging dish 24 hours before imaging. 96.2 % pure propofol was purchased from Sigma Aldrich (cat. 1572503). Propofol was serially diluted in DMEM media to prepare a 0.1% propofol solution. DMEM media was swapped with propofol media after the cells were attached to the bottom of the glass plate. Treatment groups were imaged at a time point of 30min and 24 hours.

### Fluorescence Lifetime Imaging

The NAD(P)H and FAD fluorescence lifetime images were acquired by a specially designed multi- photon microscope (Marianas, 3i), combined with a 40X water immersion objective (1.1 NA) and a tunable (680nm- 1080nm) Ti: sapphire femtosecond laser (COHERENT, chameleon Ultra II). During imaging, cell imaging dishes were placed in a stage-top incubator to maintain the environment of the cells at 37°C, 5% CO_2,_ and 85% relative humidity. NAD(P)H fluorescence was excited at 750nm with a laser power of 10-15mW, and FAD fluorescence was excited at 890nm with a laser power of 13-25mW at the specimen level. NAD(P)H and FAD FLIM images were obtained consecutively by photomultiplier tube (PMT) detectors attached to a time-correlated single-photon counting (TCSPC) electronics module (SPC 150N, Becker & Hickl). Each FLIM image was 256×256 pixels in size, acquired with a pixel dwell time of 50µs and a 3 frame repeats. The collection time was 40s and 60s for NAD(P)H and FAD, respectively. FLIM images of both NAD(P)H and FAD were captured in at least six random locations for each imaging dish. Results include three technical replicates to ensure the reliability of the results. For the instrument response function, a second harmonic generated signal of urea crystal was excited at 900m and measured with the NAD(P)H channel.

### Cell-level image analysis

SPCm was used to convert spc Images to sdt images. After that, Fluorescence lifetime decays were analyzed by SPCImage (Becker & Hickel). The decay curve of each pixel was processed with the measured instrument response function (IRF), which was obtained by the second harmonic generation of urea crystal, and then fitted into a two-exponential model. The weight-average fluorescence lifetime was calculated for each pixel with MATLAB to obtain corresponding lifetime images.

Cell segmentation was performed by using cellpose to extract fluorescence lifetime features from each cell. About 500-700 cells were extracted for each group with each experimental repeat. Finally, image processing was performed using a customized script in MATLAB and the pixel-averaged values for eight NADH(P)H τ_1_, NAD(P)H τ_2_, NAD(P)H τ_m_, NAD(P)H α_1_, FAD τ_1_, FAD τ_2_, FAD τ_m_, and FAD α_1_ were acquired for each cell.

### Osmolarity test

Osmolarity differences between the propofol-infused DMEM media and DMEM culture media (without propofol) were checked using the Precision Systems Osmette™ Micro-Osmette Osmometer. Results are summarized in Table 1.

**Table 1:** Osmolarity results of DMEM control media and DMEM media with 0.1% Propofol.

| Solutions | Osmolarity (mOsm/L)<br>Trial-1 | Osmolarity (mOsm/L)<br>Trial-2 | Mean |
| --- | --- | --- | --- |
| DMEM | 344 | 339 | 346.33 |
|  | 342 | 348 |  |
|  | 347 | 358 |  |
| 0.1% Propofol | 335 | 334 | 336 |
|  | 332 | 343 |  |
|  | 332 | 340 |  |

### Statistical analysis

Statistical analysis was performed using the data analysis and graphing software Origin Pro 2025. We performed a paired t-test to check the variances between the control, 30min and 24-hour treatment groups and to determine the statistical significance. A *p-value* of 0.05 was set to indicate significance.

### Results and Discussion

Representative FLIM images of propofol-treated MDA-MB-231 cells indicated significant morphological changes, specifically cell shrinkage, after 30min and 24hr of propofol treatment (Fig. 1). Changes in cell size could arise from osmotic differences between the control and propofol media conditions. Therefore, a comparative osmolarity assessment was performed between the DMEM culture media and propofol solution. The osmolarity results validated that there were no significant osmotic variations between the two solutions (Table 1).

**Figure 1:**
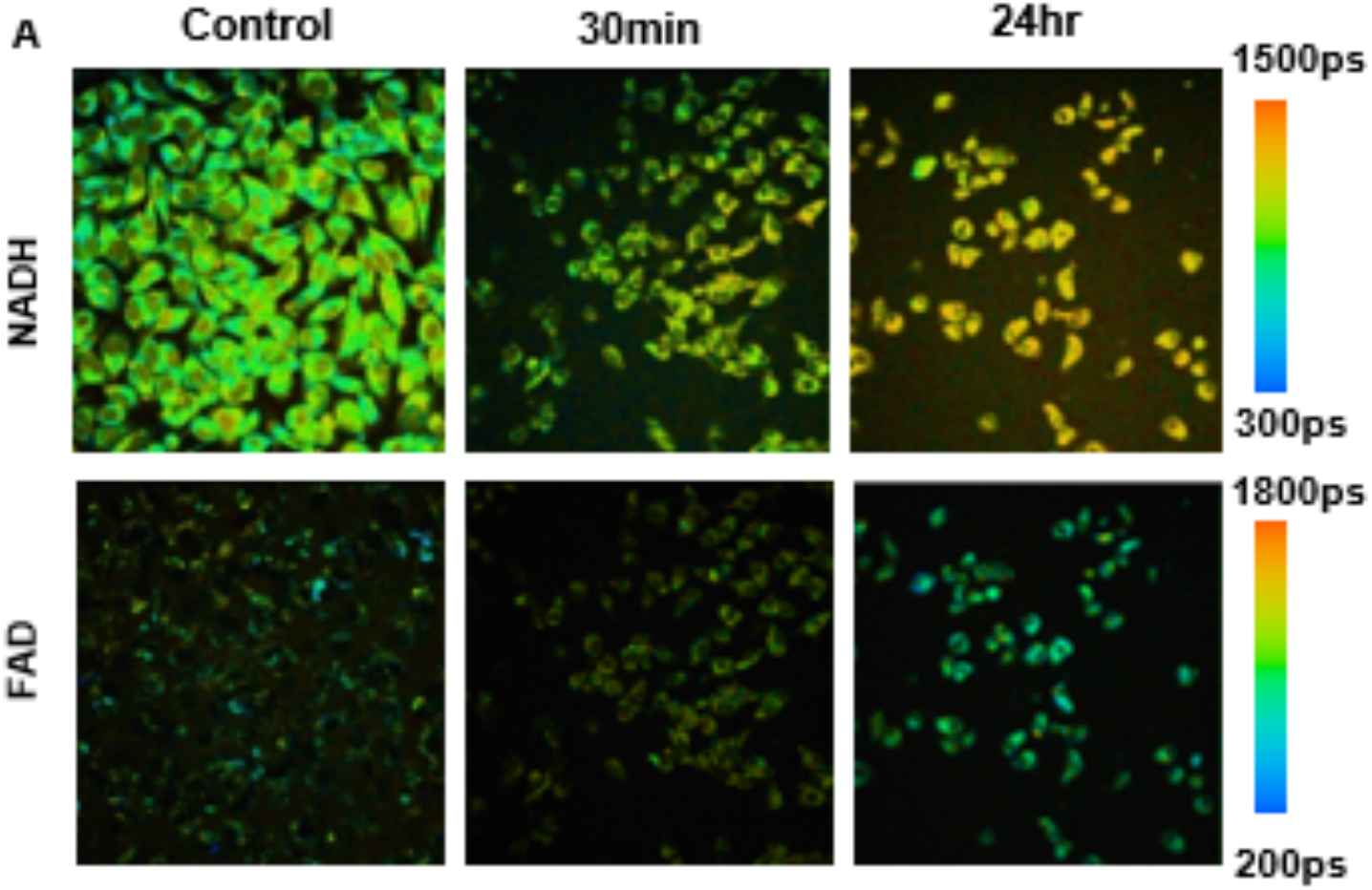
Representative FLIM images of control and 0.1% propofol-treated MDA-MB-231 cells.

The FLIM analysis indicates a shift in the mean fluorescence lifetimes of NADH and FAD signals for MDA-MB 231 cells treated for 30 minutes and 24 hours with 0.1% propofol (Fig. 2-3). From the analysis, we observed a significant decrease in the free NAD(P)H fluorescence lifetime (τ_1_) (Figure 2C) along with a decrease in the bound NAD(P)H lifetime (τ_2_) (Fig 1D) compared to the MDA-MB- 231 control groups. This decrease led to an ultimate decrease in the mean lifetime of NAD(P)H (τ_m_) (Fig 2A).

**Figure 2:**
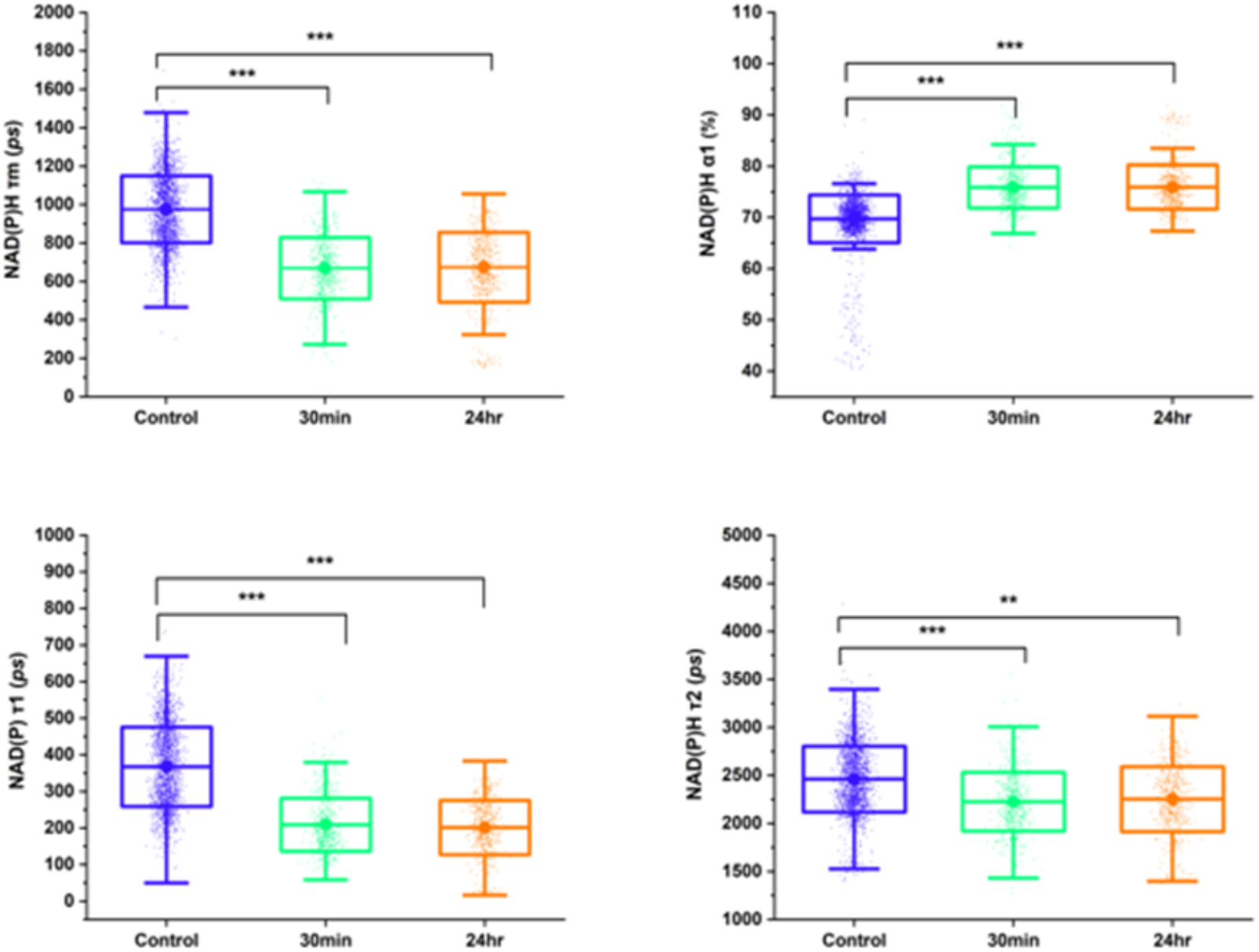
(A) Mean fluorescence lifetime of NAD(P)H is reduced in MDA-MB-231 cells with 30 minutes and 24 hours of propofol treatment. (B) Fraction of free NAD(P)H, α_1_, is increased with 30 minutes and 24 hours of propofol treatment in MDA-MD-231 cells. (C) NAD(P)H short lifetime, τ_1_, and NAD(P)H long lifetime, τ_2_, are decreased with propofol treatment. Data is presented as a boxplot with mean ± standard deviation. Each data point is an individual cell, n=3 experimental repeats. Results were considered statistically significant for *\*p< 0*.*05, **p<0*.*01, ***p<0*.*001*.

For FAD lifetime, there is a decrease in FAD short lifetime (τ_1_) (Fig. 3C), and long lifetime (τ_2_) (Fig 3D) and mean lifetime (τ_m_) (Fig 3A) for 30min propofol treatment with a slight increase in the corresponding lifetimes for 24 hr propofol treatment. We observed an increase in the FAD α_1_ or fraction of bound FAD (Fig 3B).

**Figure 3:**
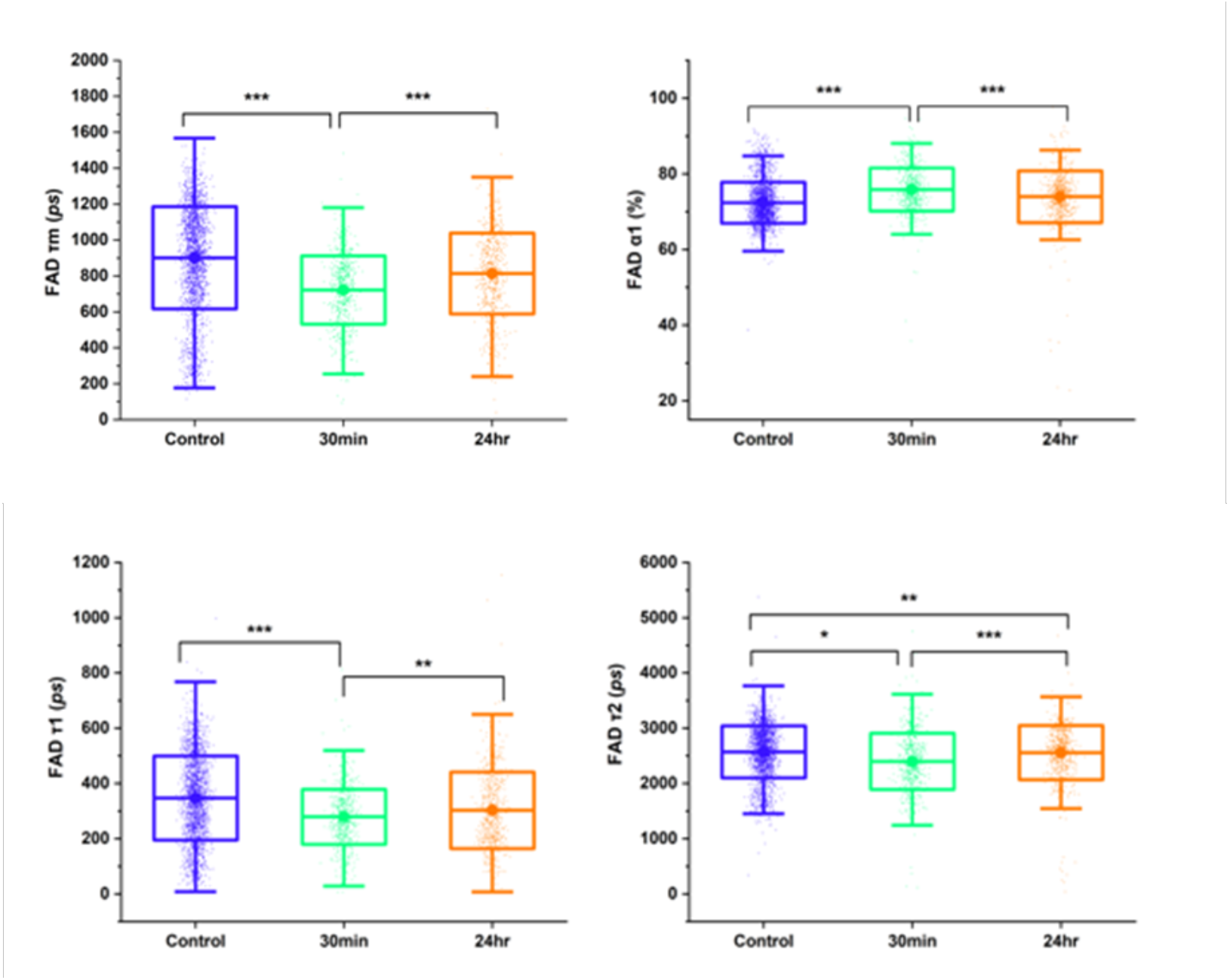
(A) Mean fluorescence lifetime of FAD is reduced in MDA-MB-231 cells with 30 minutes and 24 hours of propofol treatment. (B) Fraction of bound FAD, α_1_, is increased with 30 minutes and 24 hours of propofol treatment in MDA-MD-231 cells. (C) FAD short lifetime, τ_1_, and FAD long lifetime, τ_2_, are decreased with propofol treatment. Data is presented as a boxplot with mean ± standard deviation. Each data point is an individual cell, n=3 experimental repeats. Results were considered statistically significant for *\*p< 0*.*05, **p<0*.*01, ***p<0*.*001*.

In addition to the fluorescence lifetime changes of NAD(P)H and FAD, the fluorescence intensity values were analyzed to determine propofol effects on the optical redox ratio. The optical redox ratio, FAD/(NAD(P)H+FAD), was significantly decreased in MDA-MB-231 cells with 30 minutes and 24 hours of 0.1% propofol (Fig 4).

**Figure 4:**
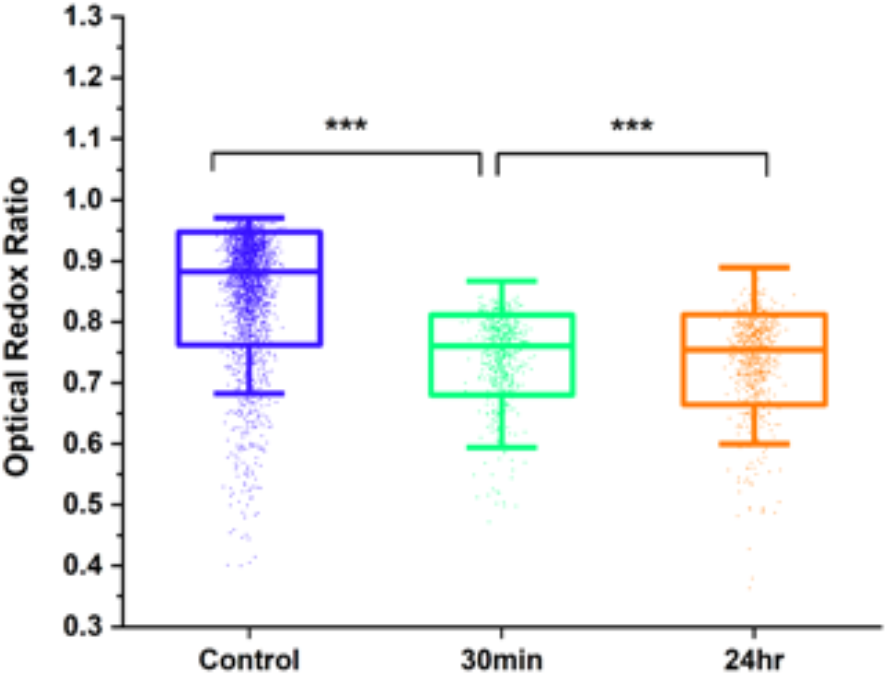
Optical redox ratio (FAD/(NAD(P)H+FAD) decreases in MDA-MD-231 cells with 30 minutes and 24 hours of 0.1% propofol treatment. Data is presented as a boxplot with median and interquartile ranges demarcated. Each data point is an individual cell, n=3 experimental repeats. Results were considered statistically significant for *\*p< 0*.*05, **p<0*.*01, ***p<0*.*001*.

The decrease in optical redox ratio suggests that propofol induces a more reduced state of the cell or a switch oxidative metabolism to glycolysis. Similarly, the observed decreases in the mean lifetimes of NAD(P)H and FAD and increases in the α1 proportion reveal that propofol treatment significantly alters the metabolism of MDA-MB-231. Altogether these results suggest autofluorescence lifetime imaging of NADH and FAD has the potential to investigate anesthesia drug effects on cellular metabolism.

## Acknowledgment

This work was funded by NIH NIGMS R35 GM142990.

## Notes

### Competing Interest Statement

The authors have declared no competing interest.

